# Seiðr: Efficient Calculation of Robust Ensemble Gene Networks

**DOI:** 10.1101/250696

**Authors:** Bastian Schiffthaler, Elena van Zalen, Alonso R. Serrano, Nathaniel R. Street, Nicolas Delhomme

## Abstract

Gene regulatory and gene co-expression networks are powerful research tools for identifying biological signal within high-dimensional gene expression data. In recent years, research has focused on addressing shortcomings of these techniques with regard to the low signal-to-noise ratio, non-linear interactions and dataset dependent biases of published methods. Furthermore, it has been shown that aggregating networks from multiple methods provides improved results. Despite this, few usable and scalable software tools have been implemented to perform such best-practice analyses. Here, we present Seidr (stylized Seiðr), a software toolkit designed to assist scientists in gene regulatory and gene co-expression network inference. Seidr creates community networks to reduce algorithmic bias and utilizes noise corrected network backboning to prune noisy edges in the networks.

Using benchmarks in real-world conditions across three eukaryotic model organisms, *Saccharomyces cerevisiae*, *Drosophila melanogaster*, and *Arabidopsis thaliana*, we show that individual algorithms are biased toward functional evidence for certain gene-gene interactions. We further demonstrate that the community network is less biased, providing robust performance across different standards and comparisons for the model organisms.

Finally, we apply Seidr to a network of drought stress in Norway spruce (*Picea abies* (L.) H. Krast) as an example application in a non-model species. We demonstrate the use of a network inferred using Seidr for identifying key components, communities and suggesting gene function for non-annotated genes.

## Introduction

The increasing accessibility of RNA sequencing in recent years has popularized large scale applications of computational biology. Two such methods, gene regulatory network (GRN) and gene co-expression network (GCN) inference, derive network structures of either transcription factor (TF) to target, or generic gene-gene associations from gene expression data. The primary goal of a GRN studies is the discovery of new regulatory interactions of known transcription factors, often in specific conditions or during certain developmental stages, or the identification of high-impact candidate genes for biotechnology applications and genome engineering.^1,2^. Conversely, GCN analysis assists researchers in functional annotation and gene-phenotype association studies ^3^.

The area of computational inference of GRNs and GCNs receives considerable attention from the scientific community with new software published regularly, but the challenging nature of the problem makes progress in the field incremental. Gene networks tend to suffer from two main issues: Firstly, the inference algorithm is often biased toward (or against) some types of regulatory interactions^4^; Secondly, the experimental limitations in the number of samples and precision of sampling often lead to low signal-to-noise ratios. In order to combat bias, Marbach *et al.* [4] proposed a voting-based scheme of a “crowd” of networks (hereinafter referred to as a community), which increased the robustness of the final aggregate network both on simulated and real data. To combat the low signal-to-noise ratio in real world networks, Coscia *et al.* [5] recently proposed a network “backboning” strategy that employs a Bayesian framework to filter non-essential edges from a dense network.

Despite the proposed methods to combat bias and noise, few studies in the field make use of either of them, often relying on a single method and a naïve edge threshold, where all edges below an often arbitrary cutoff are filtered. Some software implementations have been previously published^4,6,7^, but their applicability is narrow due to the integration of poorly optimized code. In most meta-network applications the source code of the original algorithm publications is integrated as-is, which is often coded as a proof-of-concept rather than optimized for high-throughput analysis. This severely limits the scope of the inferred network, allowing only the inference of interactions between known transcription factors and putative target genes as opposed to comprehensive all-vs-all analyses, for example. We have developed “Seidr” (stylized “Seiðr”), a feature rich software package that currently implements ensemble network inference, aggregation, backboning as well as numerous tools to interact with Seidr networks. Seidr is written in C++, can scale to tens of thousands of genes and can be executed on high performance compute clusters using message parsing interface (MPI) distributed computing, making it applicable for high-dimensional analyses, including large single-cell datasets.

We show that Seidr produces robust results in three networks produced from real-world RNA sequencing data of three model eukaryotic species. We chose to utilize real-world data as benchmarks for GRN and GCN inference software, as *in silico* gold standards are not always reliable predictors for performance on real world data, due to them being inherently subject to assumptions about the dynamics between expression quantification and gene interaction, or relying only on evidence from simpler prokaryotic species.

Finally, to demonstrate a typical use-case, we apply Seidr to a study of drought stress in Norway spruce (*Picea abies* (L.) H. Krast).

## Results

### Seidr workflow

Currently, thirteen gene network inference methods are implemented in Seidr, in three broad inference groups: correlation, mututal information (MI), and regression. Correlation based methods include Pearson correlation, Spearman correlation, topological overlap^8^ and a shrinkage estimate to partial correlation as implemented in Schäfer *et al.* [9]. In the MI group Seidr supports calculating raw MI scores using B-splines^10^, which can be further post-processed using the CLR^11^, or ARACNe^12^ algorithms. The fourth MI method implements Narromi^13^. In the regression group, Seidr provides GENIE3^14^, TIGRESS^15^, linear SVM and Elastic Net ensembles^16^, PLSNET^17^, and the topological overlap metric from WGCNA^8^. A typical Seidr workflow involves three steps (a representation can be seen in Figure 1):

1. **Inference**: Construct any number of independent gene networks using any combination of the above algorithms. This can be done for a full all-vs-all gene network or in a targeted approach (e.g., only transcription factors).
2. **Aggregation**: The group of inferred networks is aggregated into a community network using one of the supported aggregation methods.
3. **Filtering**: As aggregation usually outputs a fully-connected network, it is desirable to cut low confidence edges. Seidr can be used to estimate a hard threshold, using network scale-free fit and transitivity as lead statistics, or by applying a dynamic cutoff as suggested in Coscia *et al.* [5].
4. **Downstream analysis**: A dense or pruned network can then be utilized in a number of graph based analyses. Common choices include graph partitioning, centrality analysis or neighbourhood analyses in order to study genes (or groups of genes) related to a function of interest.

**Figure 1:**
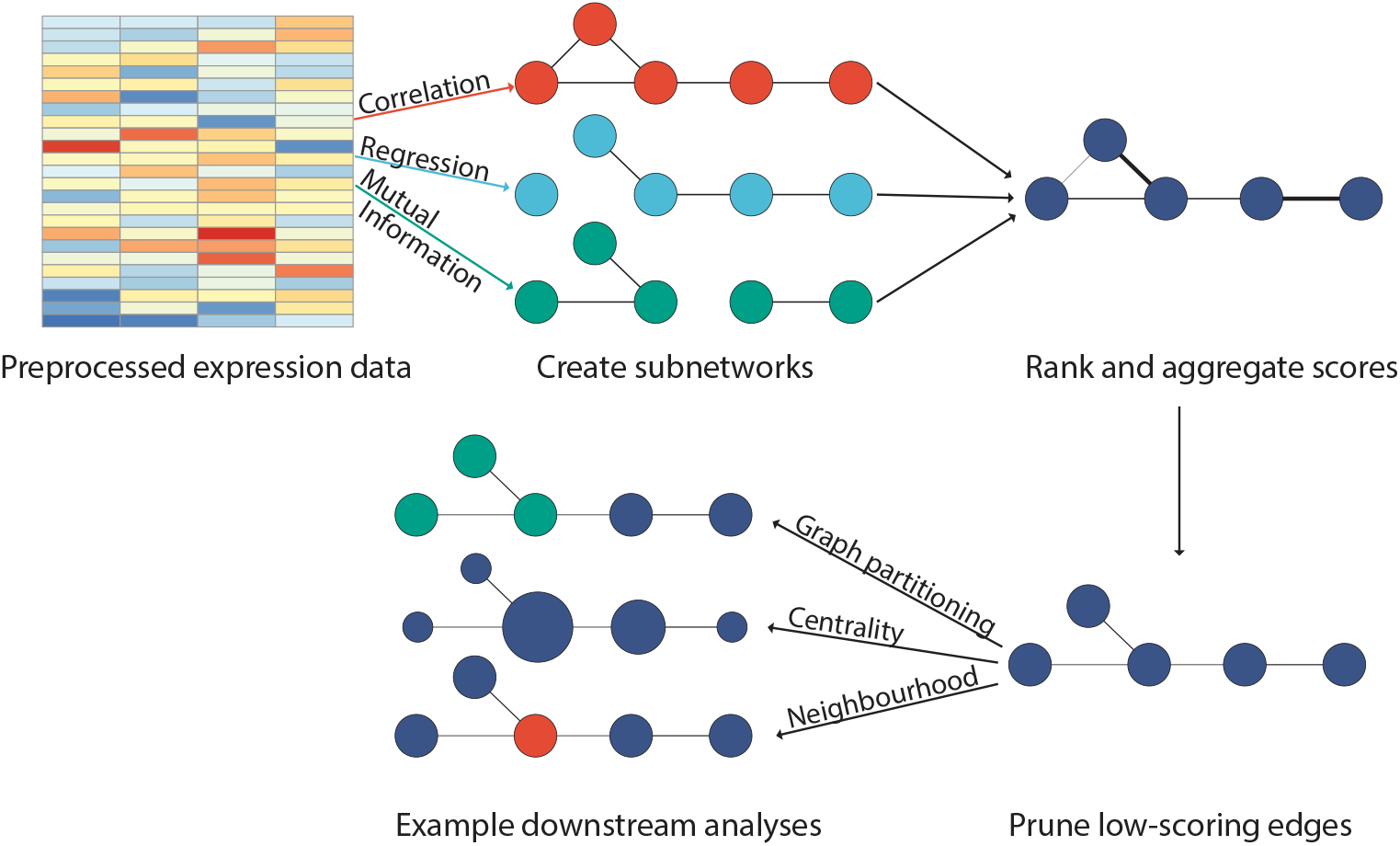
Typical Seidr pipeline. Pre-processed expression data is input into any number of network inference programs, *e. g.*, those that are implemented in Seidr itself. These networks are then rank-aggregated into a community network using a supported aggregation scheme. Networks can optionally be pruned via noise-corrected dynamic backboning or naïvely by a score cutoff. Finally, numerous downstream procedures can be used to analyse the resulting networks.

### Benchmark data and gold standards

In order to assess the performance of algorithms on real-world data we selected publicly available RNA-Seq data from the National Center for Biotechnology Information (NCBI) Sequence Read Archive (SRA) for three model species: *S. cerevisiae* (N=1399), *D. melanogaster* (N=859) and *A. thaliana* (N=1216). We selected the Biological General Repository for Interaction Datasets (BioGRID) interaction dataset for each species as a measure of ground truth for general connectivity^18^. For functional proximity, we used the Kyoto Encyclopedia of Genes and Genomes (KEGG) database as a proxy, defining an edge between two genes if they co-occur in a pathway^19^. For *S. cerevisiae*, we chose the Yeast Search for Transcriptional Regulators And Consensus Tracking (YEASTRACT) database as a measure of regulatory proximity^20^. In order to verify the predicted networks we assessed whether individual or community methods predicted higher-scoring edges for in-groups compared to out-groups. For KEGG, we tested the following null hypothesis: Links between genes annotated to the same pathway do not receive higher weights compared to genes annotated to different pathways. For YEASTRACT and and BioGRID, we formulated the null hypothesis as follows: links present in the gold standard - treated as undirected - do not receive higher weights compared to links of the same genes to those outside of the gold standard. One-sided Kruskal-Wallis tests showed strong associations for all single methods as well as the community (supplementary data 2).

### Network pruning improves sensitivity and specificity

Many gene network inference algorithms produce dense networks, meaning every gene is connected to every other gene with a calculated edge weight. To prune these networks, researchers often perform naïve pruning, which cuts all edges below an arbitrary threshold. Coscia *et al.* [5] argued that this method fails to properly take local edge distributions into account, and therefore fails to separate signal from noise accurately. We implemented their algorithm in Seidr, and performed backboning for all *δ* (the standard deviation of the expected variance of edge weights connected to a single node) in range [0.1, 0.2*, ...,* 3.5] for all algorithms implemented in Seidr and all benchmark networks. We further generated naïvely pruned networks with approximately equal edge counts to their backbone filtered counterparts. We then used all standards to generate receiver operator characteristic (ROC) and precision-recall (PR) curves for all aforementioned combinations and calculated the area under the ROC-curve (AUC) and under the PR-curve (AUPR). Briefly, the AUC reports whether a predictor ranks true positives higher than false positives and the AUPR shows how relevant the edges it ranks highly are. An *AUC* = 1 indicates a perfect predictor and *AUC* = 0.5 indicates a random predictor. We only considered networks with greater than 0.1% edge density (*i.e.* the fraction of edges left in the network compared to all theoretical edges if all genes were connected to each other), as otherwise the number of gold standard edges would be too sparse to generate ROC curves.

In general, all networks benefited from filtering edges using either method. Most networks improved by lenient backboning, pruning edges below a *δ* of 2, at which the benefit reached a plateau (see Supplementary data 2). Pruning stricter than a *δ* of 2.32 - which represents an approximate P-value of 1% - often resulted in a decrease in AUPR and a small increase in AUC, representing a trade off between sensitivity and specificity.

### Ensembles of networks improve robustness

Marbach *et al.* [4] suggested that community networks are superior to any single method due to their resilience against algorithmic bias. This idea was further tested using phosphorylation data by Hill *et al.* [21], who showed that community approaches can benefit causal networks. To understand whether different gene network inference algorithms are biased toward certain types of interactions, we created subsets of the full BioGRID standard. We split the dataset by the type of evidence as a proxy for the interaction type. Next, we created a community network for each benchmark dataset, using the inverse rank product (IRP; Zhong *et al.* [22]) algorithm, and performed the same pruning steps as for all other networks. We then recomputed the F1 scores (a measure of accuracy that combines precision and sensitivity) from the AUC and AUPR solely based on each single evidence group and compared the mean F1 of the evidence subset to the mean F1 of the full dataset. We summarized general robustness as the relative variance of mean differences to the baseline (Figure 2, top), which ranks the community network as the best cumulative score, about even with raw MI.

**Figure 2:**
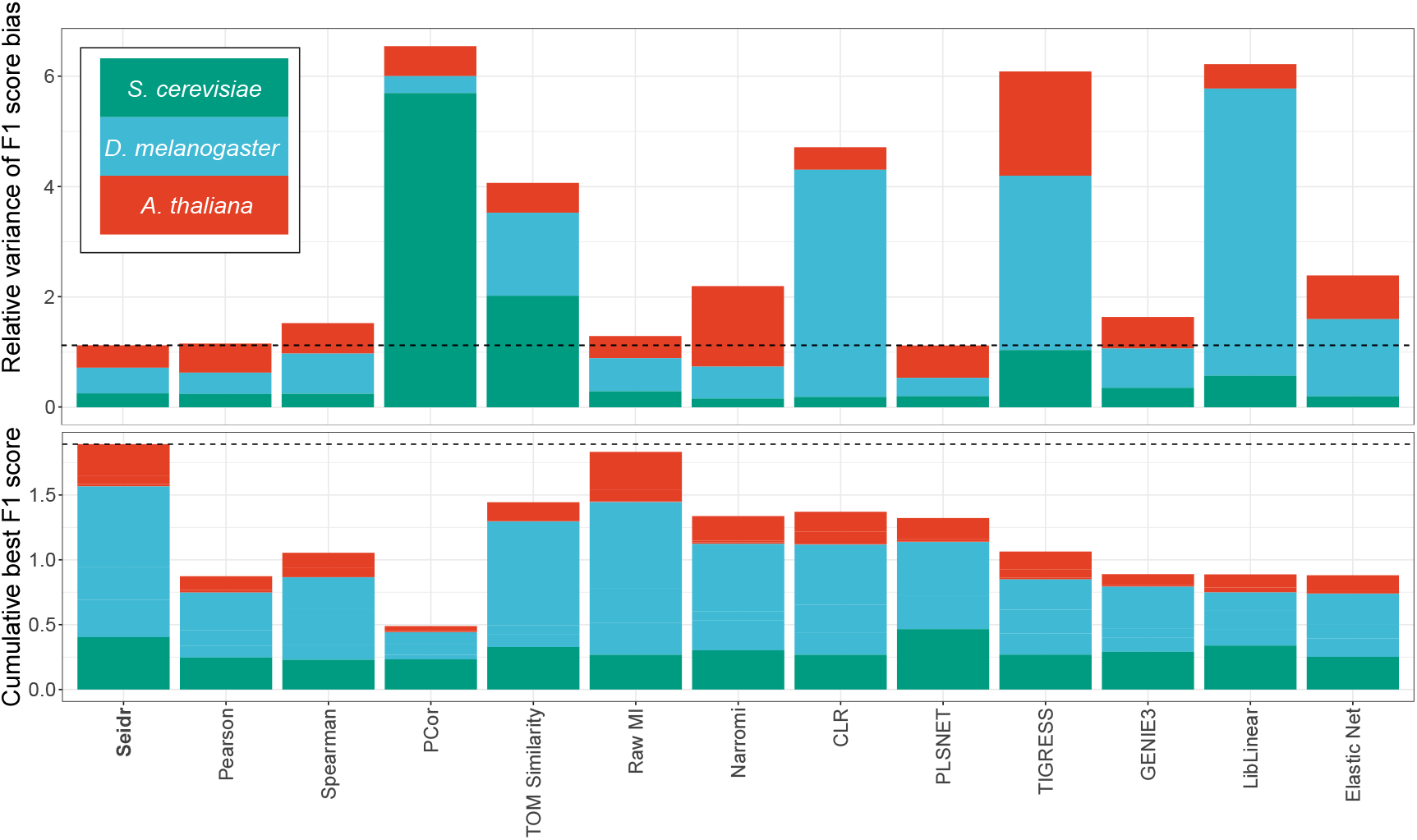
**Top:** Estimate of method bias, lower is better. Networks were evaluated on standards derived from single BioGRID evidence types and compared to the all edges from BioGRID. Increased or decreased evaluation scores (F1) suggests bias of the method toward a specific type of evidence. This plot shows the relative variance of F1 scores from subsets compared to the full dataset. **Bottom:** Cumulative method performance across all evaluations, F1 scores. For all three species as well as KEGG, BioGRID and YEASTRACT standard, the highest performing pruned network was selected for each method. The bar plot shows the sum of maximal F1 scores, shaded by species.

In order to understand whether the community network performs better than any single method, we aggregated all benchmark data and selected the highest scoring pruned network for each method. The sum of performance evaluations over all standards and species ranked Seidr as the highest scoring method, followed by raw MI (Figure 2, bottom).

### Applying Seidr to drought stress in Norway spruce

#### Drought and non-drought networks

In order to identify possible drought-specific actors in Norway spruce, we used Seidr to create a network of drought stressed needles using previously published RNA-Sequencing data^23^.

Briefly, in this experiment the authors assayed physiological and transcriptional changes in Norway spruce seedlings after inducing drought stress. Within the first five days, soil moisture was reduced from 80% field capacity (FC) to 30 % FC and subsequently held at 30% for seven days. Severe drought stress was then induced by withholding water until severe symptoms of dysfunction occurred. These conditions were held for four days after which re-irrigation started. Four days after re-irrigation started, soil moisture had returned to 80% FC. Needles were sampled at control (day zero), mild (two, four, five and 13 days) and severe (18 and 21 days) drought and after re-irrigation (25 days).

After calculating a gene network from this data, we further performed edge pruning via the backbone method, graph partitioning using InfoMap^24^ and calculated node-based centrality statistics using convenience functions implemented in Seidr. In parallel, we followed the same pipeline using a compendium of RNA-Seq data from unstressed needles as a non-specific dataset.

### Higher node centrality coincides with relevant biological functions

To investigate drought-related gene functions within the gene networks, we first used a curated list of Norway spruce orthologs of *Arabidopsis thaliana* genes with confirmed roles in drought stress^23^ (N=150, see supplementary data 4) and used Seidr-calculated node centrality metrics for each node in the stressed and un-stressed networks. We then tested whether genes in our curated dataset had significantly higher centrality values compared to all other genes or compared to a random sample of genes of the same magnitude (one-sided Kruskal-Wallis, FDR: *α <* 2.8 * 10^*−*4^). The set of curated genes was significantly higher ranked in all six centrality statistics only in the drought-specific network. While the same nodes also showed a similar trend in the non-specific network, none of the associations were statistically significant (supplementary figure 10-12). In addition, we used gene set enrichment analysis (GSEA) to test if the set of curated genes was enriched for high-centrality nodes in either network. Within the stressed network, all metrics showed a high degree of association between centrality and the set of curated genes, whereas the unstressed network only had significant enrichment in one out of six statistics (betweenness, supplementary figure 10-12).

To supplement the previous analysis with unbiased resources, we performed gene set enrichment analysis of gene ontology (GO) categories for both networks using the node centrality values as covariates (supplementary data 3). Nodes with high centrality in the stressed network were enriched for numerous response processes to biotic and abiotic stimuli, as well as metabolic processes such as nicotinamide adenine dinucleotide phosphate (NADPH) regeneration (GO:0006740), lipid biosynthesis (GO:0008610) and the mitogen-activated protein kinase (MAPK) cascade (GO:0000165). In contrast, nodes with high centrality in the unstressed network were enriched for processes such as developmental growth (GO;0048589), cell cycle (GO:0007049) and cytokinesis (GO:0000910).

#### Partitioning and Enrichment

Using the edge-pruned drought network, we partitioned the graph via InfoMap^24^ (figure 3). For each top-level module in the graph partition with more than 10 member genes (N=16), we performed gene enrichment analysis using the GO^25,26^ (supplementary data 5) and MapMan^27^ (supplementary data 6) databases as annotation. Modules 2, 5 and 6 all were significantly enriched for stress response terms (*P*_*adj*_ < 0.01). Module 2 was enriched for “defense response” (GO:0006952, *P*_*adj*_ = 2.82*e*^−11^) and “response to stimulus” (GO:0050896, *P*_*adj*_ = 2.87*e*^−3^). Module 5 was enriched for “response to oxidative stress” (GO:0006979, *P*_*adj*_ = 2.27*e*^−6^), “response to chemical” (GO:0042221, *P*_*adj*_ = 1.62*e*^−4^), and “response to redox state” (GO:0051775, *P*_*adj*_ = 5.2*e*^−4^). Finally, module 6 was enriched for “response to karrikin” - a group of plant hormones found in the smoke of burning plant material (GO:0080167, *P*_*adj*_ = 1.28*e*^−6^).

**Figure 3:**
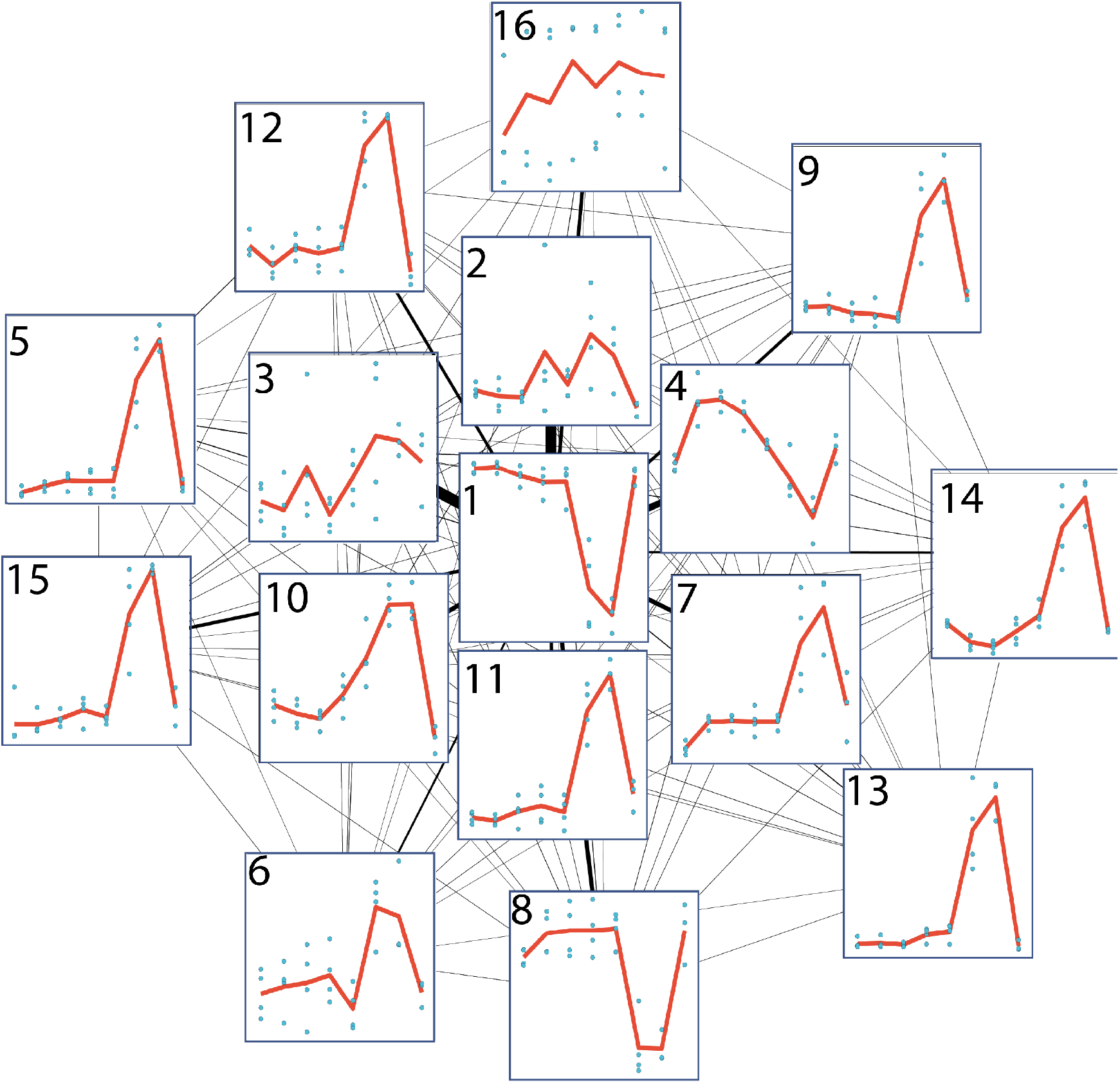
Network connectivity plot of network modules in an experiment tracking drought stress in *Picea abies*. The module number and eigengene (defined as the first principal component of the variance stabilized expression) are shown as each node, connections between nodes are represented as edges, where edge width highlights the strength of the information flow.

**Figure 4:**
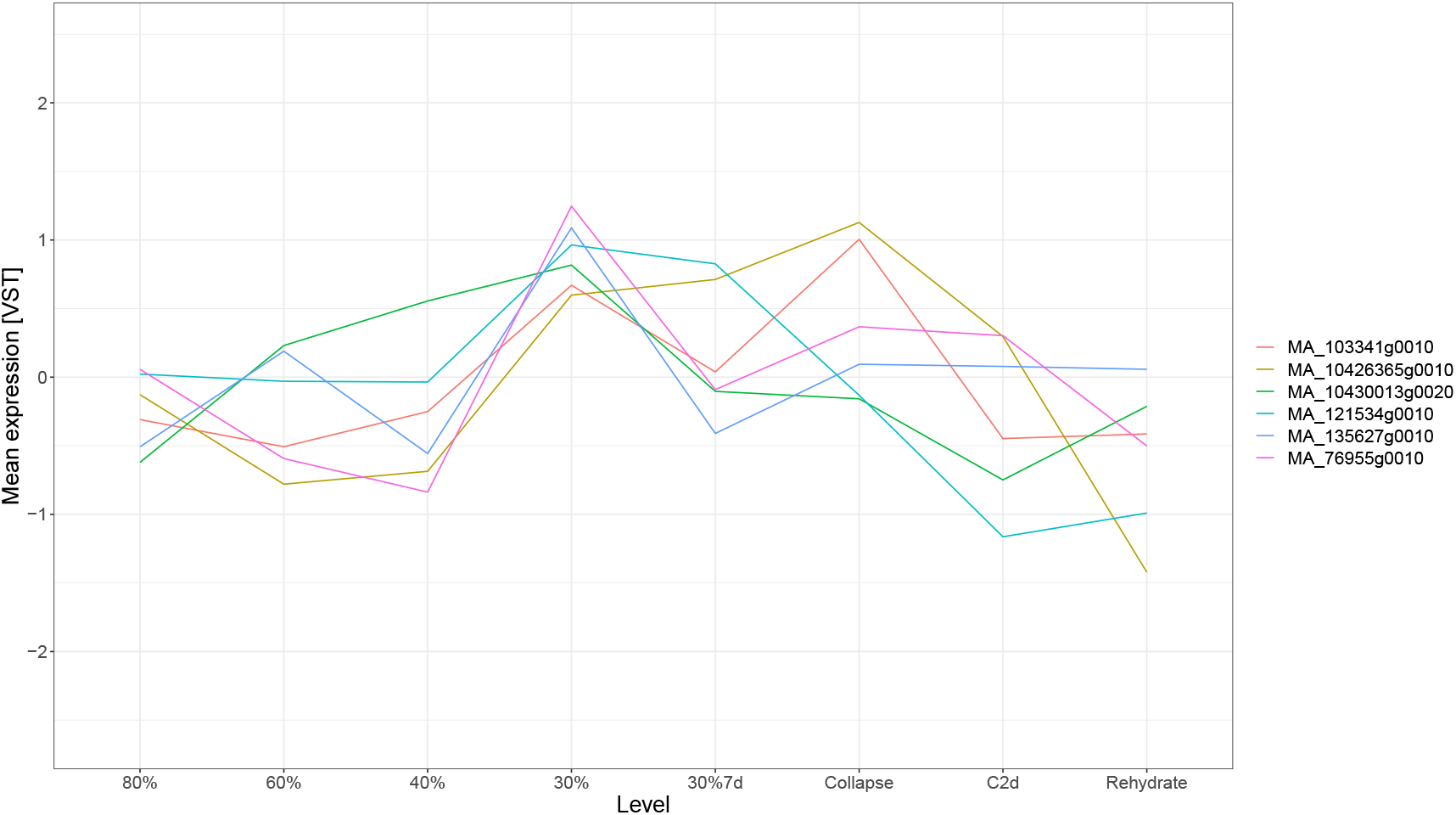
Mean expression of high-centrality transcription factors identified in module 2. Expression values are variance stabilized counts, biological replicates were summarized by their mean value. The x-axis represents the soil hydration. Data points “30%7d”, “Collapse”, “C2d” and “Rehydrate” represent seven days of continuous drought at 30% hydration, collapse in function of photosynthesis and transpiration (indicating severe dysfunction), continued collapse after two days and finally re-hydration to 80%.

#### High-centrality transcription factor analysis

In order to find possible targets for genetic modification, we calculated median-rank centrality values for all annotated transcription factors (TFs) in the network and selected the top 20 nodes. We then further narrowed down the list by only selecting transcription factors in modules 2, 5 or 6 given the previous analysis related to GO enrichment in these modules. None of the TFs were members of modules 5 and 6, but a total of six (30%) were annotated to module 2. All of these show a characteristic expression peak at 30% hydration, with “MA_103341g0010” (a NAC - NAM, ATAF1/2 and CUC2 - domain-containing 35-like TF) and “MA_10426365g0010” (an NAC domain-containing 86-like) peaking again during plant collapse. None of the TFs in module 2 were present in the curated list of genes involved in drought response.

#### Local Neighbourhood and Functional Annotation Inference

In order to assess functional inference power in the stressed network we selected three non-annotated genes with high median-rank centrality. In their local neighborhood we then selected nodes directly connected to the non-annotated gene. Gene **MA_10432675g0010** contains a cellulase (glycosyl hydrolase family 5) domain. Local neighborhood connects it to MA_10425958g0010 (Kinesin, mitochondrial isoform X1), MA_4391326g0010 (another cellulase) and MA_84105g0010, a glycoside hydrolase.

Gene **MA_69177g0010** has two non-annotated neighbours - MA_136426g0010 and MA_894367g0010 - as well as MA_41206g0010 which is annotated as a mitogen-activated kinase 3.

Gene **MA_10192193g0010**. This gene has two neighbours. MA_10192193g0020, annotated as zinc finger CONSTANS-LIKE 16, and MA_10226519g0010, which is a DNA photolyase.

## Discussion

Gene network inference, be it regulatory or co-expression networks, is an important and widely used bioinformatics tool for modern biological research, but best practice information and implementations are lacking. This consideration is especially relevant now, as the single cell sequencing boom is generating increasingly massive-scale expression datasets that synergize well with gene network analysis given their high sample count^28^. However, care must be taken as to which methods are selected to perform the inference^29^, with several new methods proposed to specifically address network inference in single cell experiments^30–32^.

As a software implementation to these ends we present Seidr, a software toolkit that enables researchers to create community networks, perform backboning and perform a variety of other tasks related to gene networks. Our toolkit is efficient and follows evidence-based well performing workflows. Currently, Seidr performs inference using thirteen distinct methods, but a high-level implementation based around multi processing (OpenMP), message parsing (MPI), and the Armadillo linear algebra framework makes the inclusion of new methods in a parallel, shared-memory architecture a straightforward effort. Further, the modular design allows users to import networks calculated with external tools into Seidr and add them to communities, or post-process them in one of many ways.

We used Seidr to reaffirm that community networks and dynamic edge pruning (backboning) both perform better than single methods, congruent with Marbach *et al.* [4] and Coscia *et al.* [5]. We show that community networks are less biased and perform very well compared to any single method in sensitivity, specificity and precision. Precision and recall are especially important in imbalanced problems such as gene regulatory networks, where true negatives far outweigh true positive labels. In addition to network aggregation, we also show that edge pruning in general, and backboning specifically, improved the characteristics of the generated networks.

To illustrate a number of applications, we used Seidr to infer a network based on a real-world study of drought response in *Picea abies*. In addition, we calculated another network for a compendium of unstressed needles.

Firstly, we calculated centrality statistics of all nodes in the drought and unstressed networks. The drought network showed more relevant associations between centrality and GO terms in our GSEA analysis. In addition, genes with putative involvement in drought stress were significantly associated with centrality values in the drought-specific network only, highlighting the importance of creating experiment specific networks.

In order to further gather context about local relationships of genes we partitioned the graph using InfoMap, which resulted in 16 distinct modules. We chose parameters that would result in more modules so as to cluster smaller, more specific groups of genes even if the expression profiles between modules were similar. Of the three modules which were enriched to stress-related GO terms (modules 2, 5 and 6), module 2 was of particular interest as the expression profile corresponded well to the abscisic acid measurements in Haas *et al.* [23]. Additionally, we identified six highly central transcription factors within that module, which present high interest targets for future genetic modification experiments.

Another application of gene networks is inference of possible gene function of non-annotated genes. Functional analysis of the neighbourhood of non-annotated genes revealed three diverse candidates, which were selected due to their high centrality values in the drought network. The first candidate MA_10432675g0010, is annotated as a cellulase. Genes in its neighborhood are similarly enzymes involved in cell wall modification. Sasidharan *et al.* [33] review various mechanisms of cell wall reorganization as a response to stresses, suggesting MA_10432675g0010 acts also in response to abiotic stresses to reorganize cell wall structure. The second candidate, MA_69177g0010, has no domain annotation. Only one of its neighbours has been annotated as a mitogen-activated kinase (MAPK). Sinha *et al.* [34] discuss the role of MAPKs in plants in cellular signaling as a response to abiotic stress, which would place MA_69177g0010 as part of this signaling chain. Finally, MA_10192193g0010 has a neighbour annotated as a photolyase and another as CONSTANS-like. Photolyases are known to deal with DNA-repair in response to light damage^35^. Drought stress can lead plants to have reduced ability to manage light and therefore generate reactive oxygen species (ROS), which can lead to DNA damage similar to excess UV-light^36,37^. The other neighbour is annotated to CONSTANS-like 16 (a gene typically associated with flowering), which has recently been shown to lead to chlorophyll accumulation when over-expressed in *Petunia*^37^. Further, a similar gene has recently been shown to improve drought tolerance in sugarcane when over-expressed^38^ via maintenance of photosynthesis and by enhancing the antioxidant and osmotic capabilities of the plant. We therefore infer MA_10192193g0010 to play a role in similar processes *i.e.* ROS detoxification and photosynthesis upkeep.

## Supporting information

Gene set enrichment results for stressed and unstressed networks

Curated and random gene lists for GSEA analyses

GO enrichment results for clusters 1-16

MapMan enrichment results for clusters 1-16

## Supplementary data

1. Dataset accessions
2. Supplementary S1
3. Gene set enrichment results for stressed and unstressed networks
4. Curated and random gene lists for GSEA analyses
5. GO enrichment results for clusters 1-16
6. MapMan enrichment results for clusters 1-16

## 1 Acknowledgments

## 2 Author contributions

BS, ND and NRS conceptualized all analyses and selected appropriate datasets for the experiments. BS wrote the majority of the software, performed benchmarking and centrality analyses, created all figures and wrote the manuscript. EZ performed clustering, enrichment and transcription factor analyses as well as curated the list of genes putatively involved in drought stress. AS contributed source code and optimizations to Seidr. BS and ND developed the inference pipeline. BS, ND and NRS edited the manuscript.

## 3 Software and Data availability

Seidr is available at https://github.com/bschiffthaler/seidr. Relevant scripts used in this publication are available at https://github.com/bschiffthaler/seidr-manuscript/. The raw network data for all networks in this publication can be downloaded at ftp://130.239.72.87/Facility/Manuscript/seidr.

## Methods

### Count data

For *A. thaliana*, 1227 accessions were collected from the NCBI SRA and quantified against the TAIR10 assembly using salmon/0.13.1 with default options. For *S. cerevisiae* 2129 accessions from the SRA were quantified using salmon/0.11.2 against the EnsEMBL (r93) assembly. For *D. melanogaster*, 1316 accessions were quantified using salmon/0.11.2 against the FlyBase 6.25 assembly. Each dataset was filtered to have at least 75% aligned reads, resulting in 1216, 1399 and 859 accessions post-filtering respectively. The count data was imported into R^39^ using the tximport package^40^, transcript counts were summarized to genes, and the variance stabilizing transformation was applied using DESeq2^41^. Genes with constant expression (zero variance) were removed. Finally, the median expression of all genes in a sample was subtracted from each gene in that sample, median-centering it. Detailed lists of all accessions can be found in supplementary data 1.

### Seidr benchmark networks

All network inference was done using seidr/0.13.1, with default options unless otherwise specified. In the el-ensemble, llr-ensemble, tigress, pearson, plsnet, and pcor subprograms, the “–scale” option was used to transform data to z-scores prior to inference. The networks were aggregated using the inverse rank product method. Each network backbone was then calculated (for all individual and the community networks) and the edges were filtered for all values of *δ* in range [0.1, 0.2*, ...,* 3.5], creating 35 increasingly stringent subsets of the network. Hard cutoffs were then used to create another 35 networks that match the backboned ones in density, i.e. have the same number of edges. Finally, only networks with at least 0.1% edge density were considered for benchmarking. Briefly, edge density is the fraction of edges in a network compared to all theoretical edges, where all genes are connected to all genes. As we only allow one edge between any two genes, this is the triangular matrix without a diagonal: 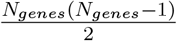. An edge density of 0.1% equates 22167 edges in *S. cerevisiae*, 94222 in *D. melanogaster*, and 485862 in *A. thaliana*.

### Norway spruce network inference

The drought network was inferred using raw data from Haas *et al.* [23], whereas the unstressed network was inferred using a collection of needles from diverse experiments at the Umeå Plant Science Center (Schneider *et al.* [42] and Nystedt *et al.* [43] as well as two unpublished datasets) and other public data (SRA accessions SRP014311, SRP093366 and SRP116733). Both networks were inferred analogous to the benchmark networks. The raw inferred networks are available at ftp://130.239.72.87/Facility/Manuscript/seidr/.

### Norway spruce network post-processing

The aggregated network was pruned at *δ* = 1.28 (approx. P-value of 0.1) using the backboning method. The resulting graph was partitioned using InfoMap^24^ version 1.2.1 with parameters:

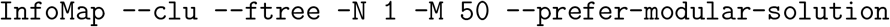

Finally, we calculated centrality metrics for nodes via PageRank^44^, Strength (weighted degree), Eigenvector centrality, Laplacian centrality^45^, Betweenness centrality and Closeness centrality using the default settings in seidr/0.14. In order to summarize all centrality values to a single value per node, we ranked nodes within each algorithm and calculated the median per-node rank across algorithms.

### Statistical analysis

Unless otherwise noted, all statistical analysis was performed in R (version 4.0.2). Gene set enrichment analysis was performed using the fgsea package (version 1.16.0)^46^. All GO enrichment was performed using in-house software implementing the parent child adjusted test from Grossmann *et al.* [47], while MapMan enrichment uses Fisher’s exact test. Both use all connected nodes in the network as the test background.

### Gold standards

All gold standard datasets were retrieved in February 2020 from their respective sources (BioGRID^18^, KEGG^19^, YEASTRACT^20^). For BioGRID, subsets of the full dataset were created based on the evidence type noted for each interaction. For KEGG, the pathway annotations were taken from the R “AnnotationDbi”^48^ package. A positive edge in the KEGG standard was defined by two genes sharing at least one pathway, a negative edge between two genes that share no pathways. The “seidr roc” subprogram was then used to calculate true positive rate, false positive rate and precision for all networks and standards. For BioGRID and YEASTRACT where there are no true negatives, we labeled any non-positive edge a negative edge. All plots were then created in R using the “ggplot2” package^49^.

## Notes

ftp://130.239.72.87/Facility/Manuscript/seidr/

### Competing Interest Statement

The authors have declared no competing interest.

