## Supplementary S1 for "Seiðr: Efficient Calculation of Robust Ensemble Gene Networks"

### Supplementary results

#### Edge weight information capture

The goal of this analysis was to check whether edge weights relate to links present in several databases (KEGG, BioGRID, YEASTRACT). For KEGG, we tested the following null hypothesis: Links between genes annotated to the same pathway do not receive higher weights compared to genes annotated to different pathways. For YEASTRACT and and BioGRID, we formulated the null hypothesis as follows: Links present in the gold standard - treated as undirected - do not receive higher weights compared to links of the same nodes to other genes. See fig 1-7.

#### Edge pruning methods

We compared naiv̈e edge pruning and back-boning with respect to the effect on AUC, AUPR and F1 scores of all benchmark networks and databases. See fig 8-9.

#### Enrichment analyses

After curating a list of genes with putative involvement in drought stress (GsOI), we tested whether these genes were associated with network centrality in the spruce drought network. Throughout the analyses, nodes from the GsOI list are strongly associated with higher centrality values. See fig. 10-12.

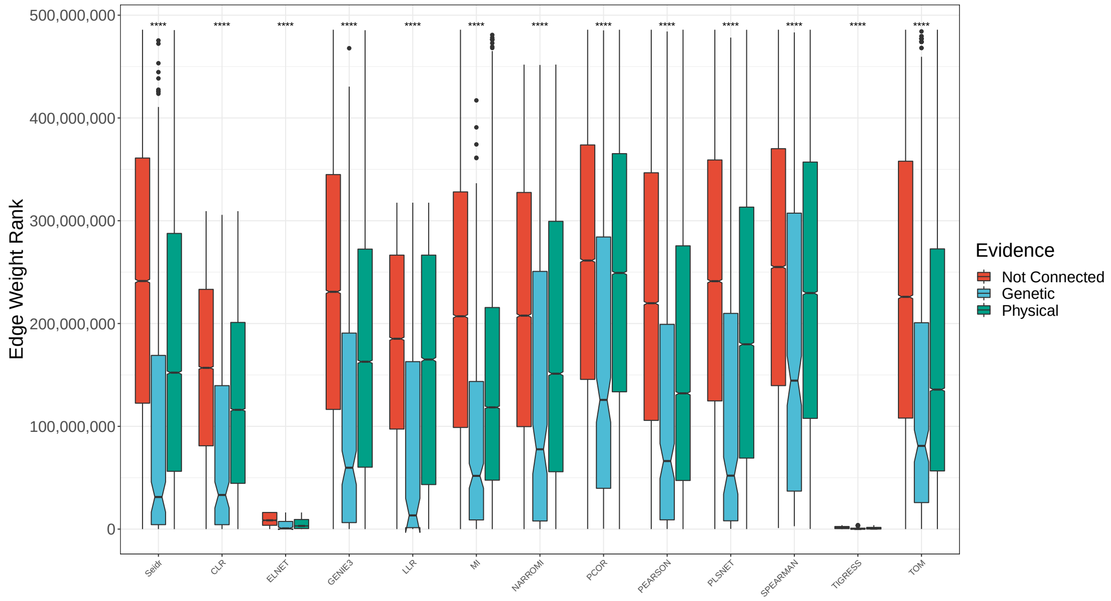


Figure 1: Box plots of edge weights (ranks) in the *Arabidopsis thaliana* network using the BioGRID database. Links shown in red do not have evidence in BioGRID, links in blue have genetic evidence, links in green have physical evidence. Lower is better. Statistical test: one-sided Kruskal wallis

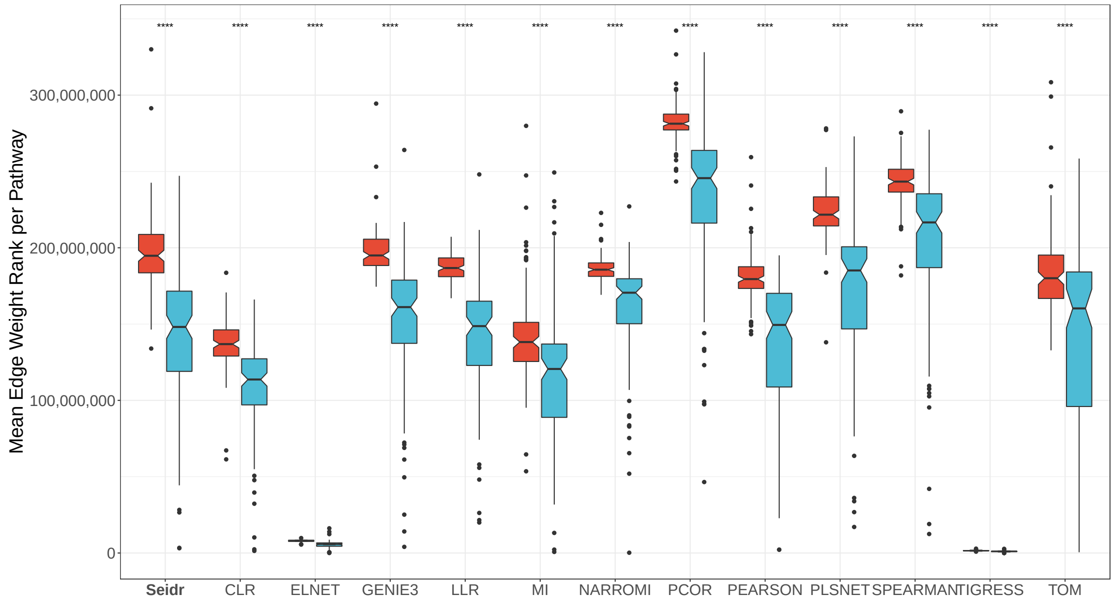


Figure 2: Box plots of edge weights (ranks) in the *Arabidopsis thaliana* network using the KEGG database. Links shown in red are not in the same KEGG pathway, links in blue are. Lower is better. Statistical test: one-sided Kruskal wallis

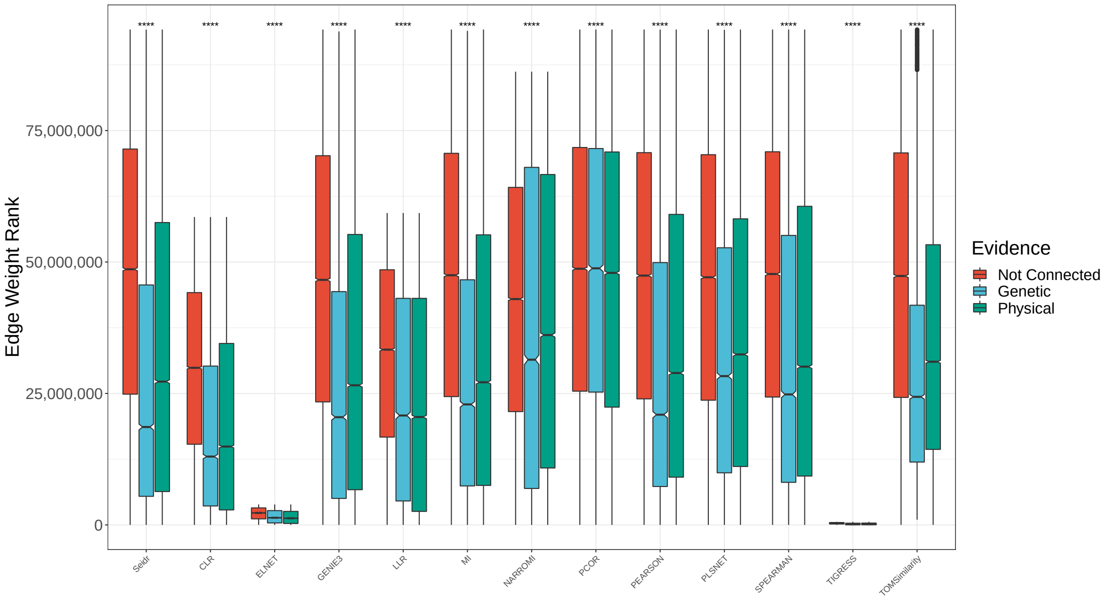


Figure 3: Box plots of edge weights (ranks) in the *Drosophila melanogaster* network using the BioGRID database. Links shown in red do not have evidence in BioGRID, links in blue have genetic evidence, links in green have physical evidence. Lower is better. Statistical test: one-sided Kruskal wallis

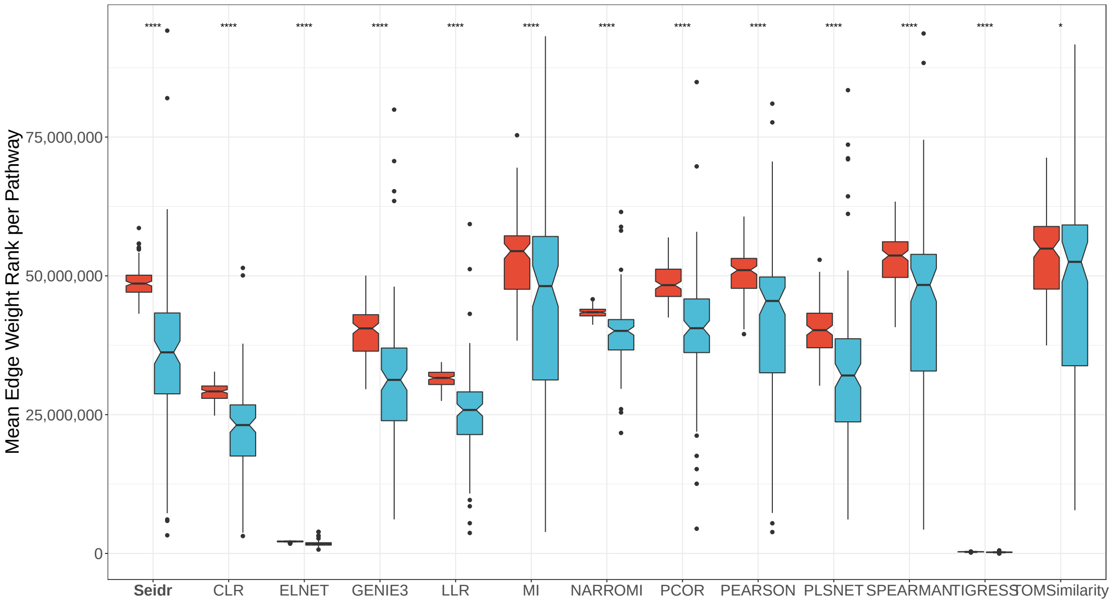


Figure 4: Box plots of edge weights (ranks) in the *Drosophila melanogaster* network using the KEGG database. Links shown in red are not in the same KEGG pathway, links in blue are. Lower is better. Statistical test: one-sided Kruskal wallis

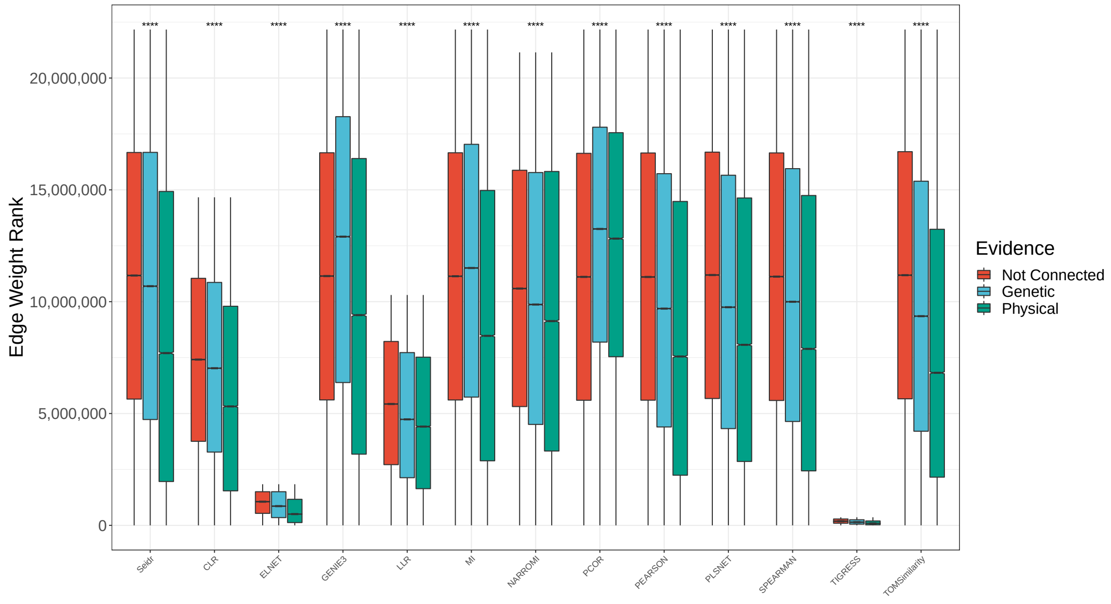


Figure 5: Box plots of edge weights (ranks) in the *Saccharomyces cerevisiae* network using the BioGRID database. Links shown in red do not have evidence in BioGRID, links in blue have genetic evidence, links in green have physical evidence. Lower is better. Statistical test: one-sided Kruskal wallis

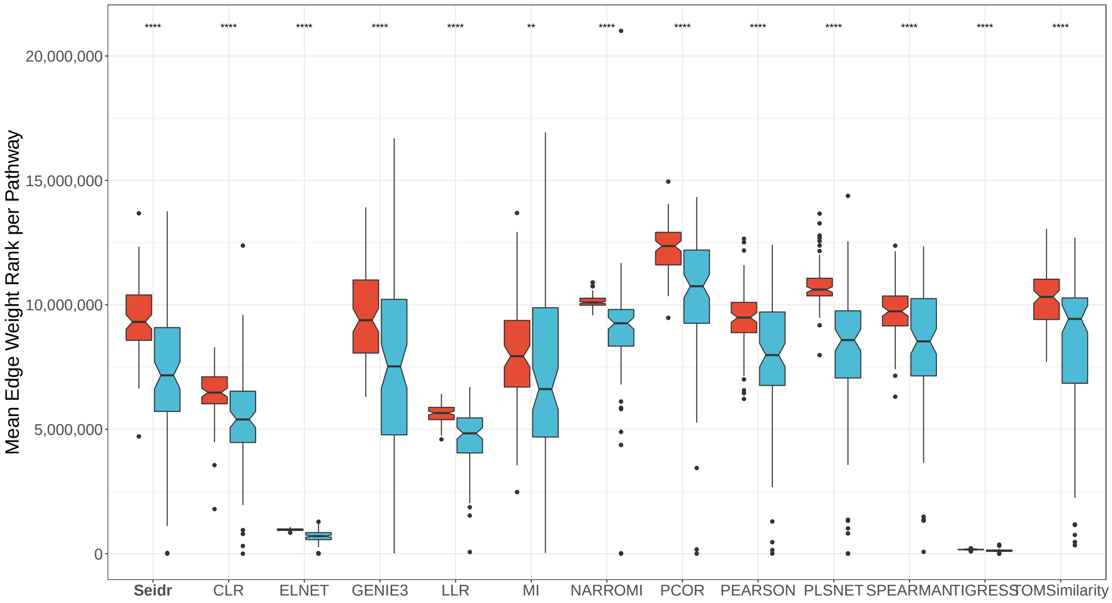


Figure 6: Box plots of edge weights (ranks) in the *Saccharomyces cerevisiae* network using the KEGG database. Links shown in red are not in the same KEGG pathway, links in blue are. Lower is better. Statistical test: one-sided Kruskal wallis

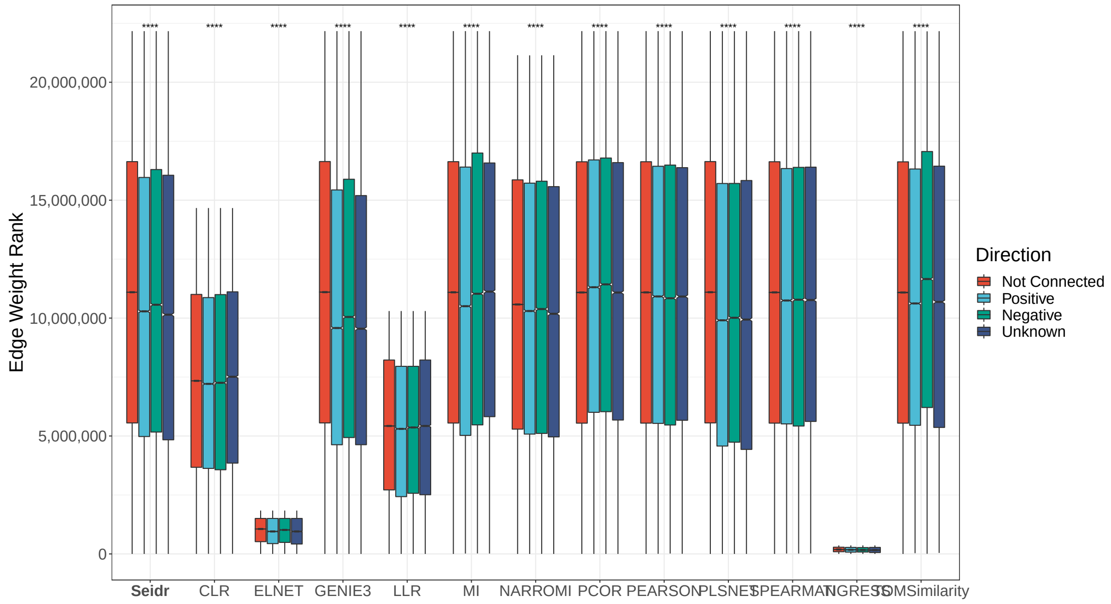


Figure 7: Box plots of edge weights (ranks) in the *Saccharomyces cerevisiae* network using the YEASTRACT database. Links shown in red are not linked in the YEASTRACT database. Links in light blue and green are positive and negative regulators respectively. Links in dark blue are associated with unknown regularity direction.

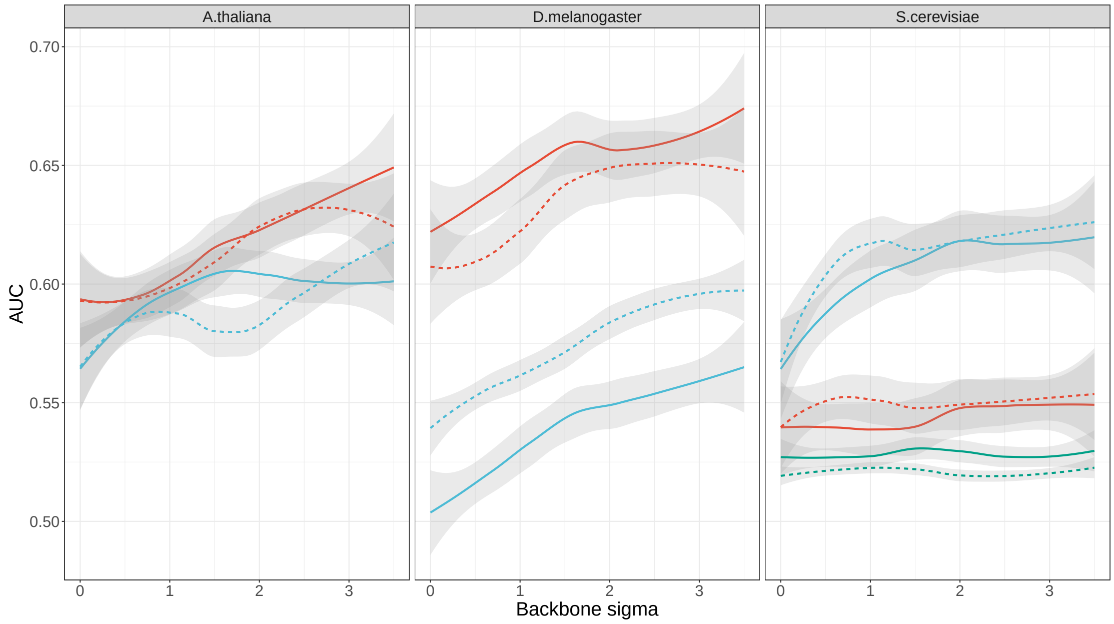


Figure 8: Comparison of naïve edge pruning (dashed lines) and back-boning (straight lines) with respect to the effect network AUC using KEGG (red), BioGRID (blue) or YEASTRACT (green) as a reference.

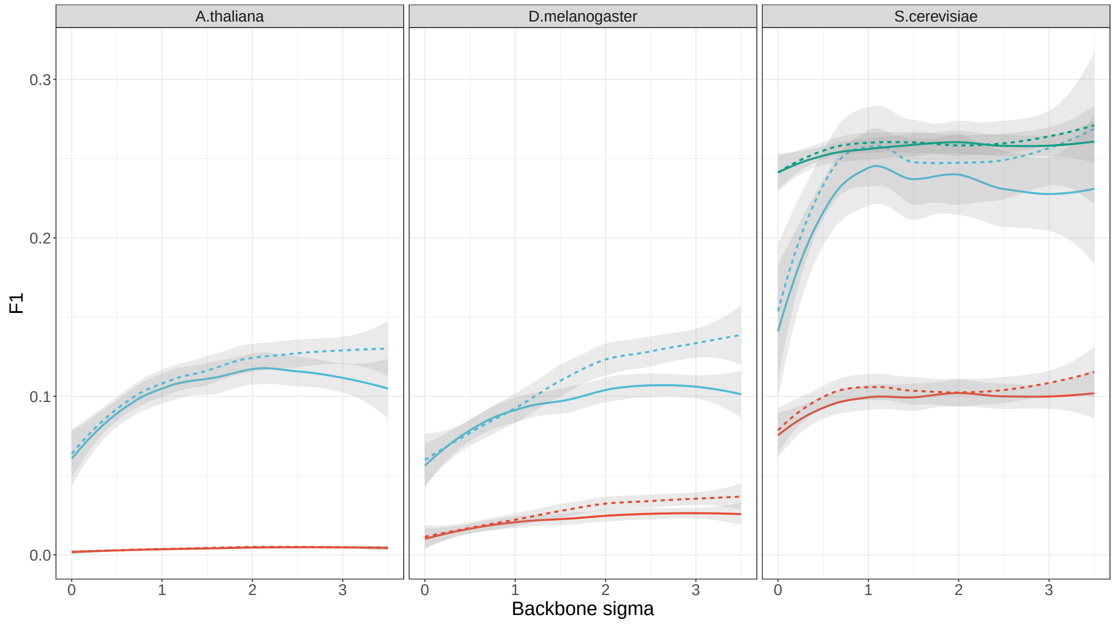


Figure 9: Comparison of naïve edge pruning (dashed lines) and back-boning (straight lines) with respect to the effect network F1 using KEGG (red), BioGRID (blue) or YEASTRACT (green) as a reference.

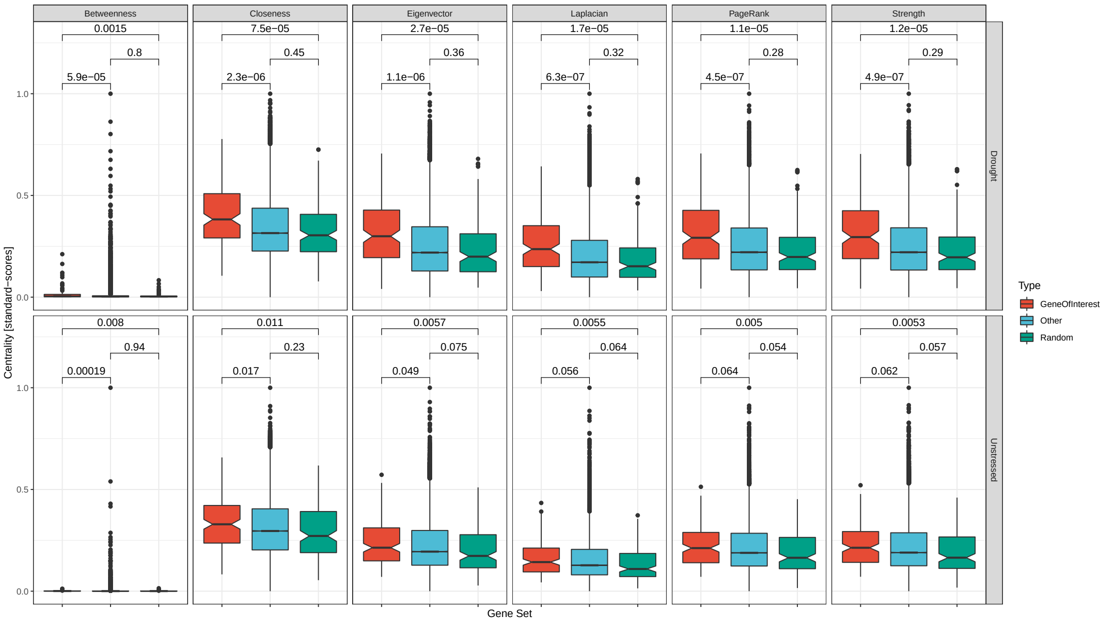


Figure 10: Comparison of GSEA analysis of three sets of genes in two networks (drought and unstressed) using median centrality as the score covariate. In red, a curated set of genes with putative involvement in drought stress. In green, a selection of random nodes from within the network with the same magnitude as the curated gene list. In blue, all other genes in the network.

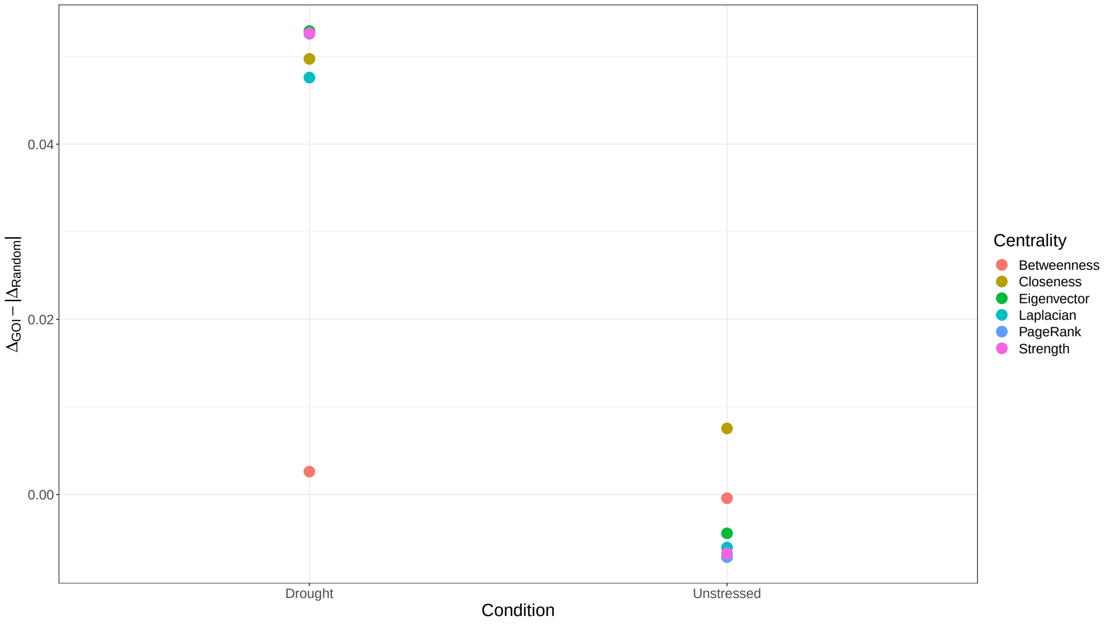


Figure 11: Comparison of GSEA analysis of two sets of genes in two networks (drought and unstressed) using median centrality as the score covariate. This plot shows the absolute difference (magnitude) of p-values between genes in the curated list and a random selection of genes of the same magnitude.

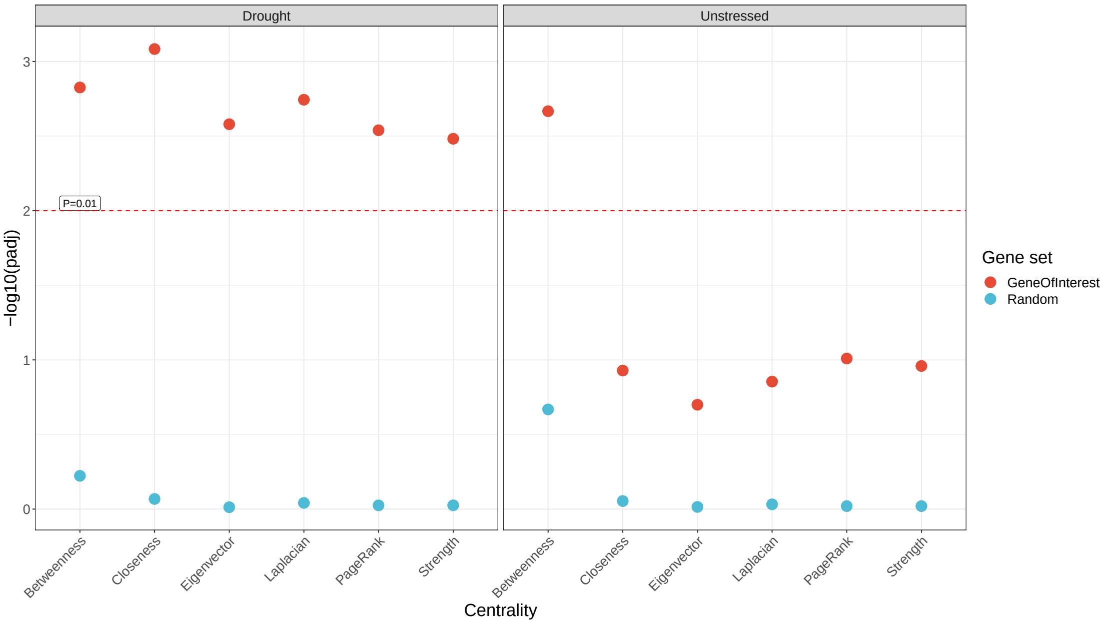


Figure 12: Comparison of GSEA analysis of two sets of genes in two networks (drought and unstressed) using median centrality as the score covariate. In red, a curated set of genes with putative involvement in drought stress. In blue, a selection of random nodes from within the network with the same magnitude as the curated gene list.
